# Ancestral SARS-CoV-2 driven antibody repertoire diversity in an unvaccinated individual correlates with expanded neutralization breadth

**DOI:** 10.1101/2022.10.19.512979

**Authors:** Suprit Deshpande, Mohammed Yousuf Ansari, Jyoti Sutar, Payel Das, Nitin Hingankar, Sohini Mukherjee, Priyanka Jayal, Savita Singh, Anbalagan Anantharaj, Janmejay Singh, Souvick Chattopadhyay, Sreevatsan Raghavan, Mudita Gosain, Supriya Chauhan, Shweta Shrivas, Chaman Prasad, Sangeeta Chauhan, Neha Sharma, Pradipta Jana, Ramachandran Thiruvengadam, Pallavi Kshetrapal, Nitya Wadhwa, Bhabatosh Das, Gaurav Batra, Guruprasad Medigeshi, Devin Sok, Shinjini Bhatnagar, Pramod Kumar Garg, Jayanta Bhattacharya

## Abstract

Understanding the quality of immune repertoire triggered during natural infection can provide vital clues that form the basis for development of humoral immune response in some individuals capable of broadly neutralizing pan SARS-CoV-2 variants. We assessed the diversity of neutralizing antibody responses developed in an unvaccinated individual infected with ancestral SARS-CoV-2 by examining the ability of the distinct B cell germline-derived monoclonal antibodies (mAbs) in neutralizing known and currently circulating Omicron variants by pseudovirus and authentic virus neutralization assays. The ability of the antibodies developed post vaccination in neutralizing Omicron variants was compared to that obtained at baseline of the same individual and to those obtained from Omicron breakthrough infected individuals by pseudovirus neutralization assay. Broadly SARS-CoV-2 neutralizing mAbs representing unique B cell lineages with non-overlapping epitope specificities isolated from a single donor varied in their ability to neutralize Omicron variants. Plasma antibodies developed post vaccination from this individual demonstrated neutralization of Omicron BA.1, BA.2 and BA.4 with increased magnitude and found to be comparable with those obtained from other vaccinated individuals who were infected with ancestral SARS-CoV-2. Development of B cell repertoire capable of producing antibodies with distinct affinity and specificities for the antigen immediately after infection capable of eliciting broadly neutralizing antibodies offers highest probability in protecting against evolving SARS-CoV-2 variants.

**Importance:** Development of robust neutralizing antibodies in SARS-CoV-2 convalescent individuals is known, however varies at population level. We isolated monoclonal antibodies from an individual infected with ancestral SARS-CoV-2 in early 2020 that not only varied in their B cell lineage origin but also varied in their capability and potency to neutralize all the known VOC and currently circulating Omicron variants. This indicated establishment of unique lineages that contributed in forming B cell repertoire in this particular individual immediately following infection giving rise to diverse antibody responses that could compensate each other in providing broadly neutralizing polyclonal antibody response. Individuals who were able to produce such potent polyclonal antibody responses after infection have a higher chance of being protected from evolving SARS-CoV-2 variants.

## Observation

Currently the COVID-19 pandemic is primarily driven by the Omicron lineage variants globally (https://nextstrain.org/ncov/gisaid/global/6m). Recently, few studies highlighted a significant reduction in effectiveness of the vaccine, therapeutic monoclonal antibodies and natural infection induced neutralizing antibody responses against the Omicron BA.1, BA.2 and rapidly emerging BA.4/BA.5 variants (1–6). Little is known about the antibody responses mounted in individuals previously infected with the ancestral SARS-CoV-2 (Wu-1/CoV2) prior to vaccination and capable of broadly neutralizing evolving VOCs including the currently circulating and rapidly emerging Omicron variants. We previously reported isolation of five RBD-specific neutralizing monoclonal antibodies from an unvaccinated Indian individual (donor C-03-0020) who was infected with ancestral SARS-CoV-2 (7). In the present study, we report isolation and characterization of two additional mAbs from the same individual: THSC20.HVTR11 and THSC20.HVTR55 (Fig. S1) by RBD-specific single B cell sorting and following same pipeline reported before (7) that differed in their B cell origin. These two new mAbs varied in their ability to neutralize Omicron variants as shown in Fig.S1A. While THSC20.HVTR11 was found to neutralize Omicron BA.1 and BA.2 potently but failed to neutralize Delta (B.1.617.2) and Kappa (B.1.617.1), THSC20.HVTR55 failed to neutralize all Omicron variants (Table S1). The loss of the ability of THSC20.HVTR11 to neutralize Delta and Kappa was found to be correlated with the presence of L452R mutation (Fig.S1B). This could also likely be the reason for the inability of THSC20.HVTR11 to neutralize Omicron BA.4/BA.5. Both the newly isolated mAbs (THSC20.HVTR11 and THSC20.HVTR55) not only demonstrated strong binding affinity but also differed in their epitope specificities on RBD (Fig. S1 C-E). Remarkably, all the seven mAbs isolated from this donor (Fig.1A) were found to be derived from distinct B cell germlines (IGVH3-30, IGVH7-4-1, IGVH1-69, IGVH3-53, IGVH4-39, IGVH5-51, IGVH1-18 linked to the variable heavy chain IgG sequences and IGVL2-11, IGVL3-1, IGVL1-40, IGVL2-23, IGVL3-21, IGVK1-5, IGVK3-20 linked to the variable light chain IgG sequences respectively), when compared with those found in the CoVAbDAb database (8) (Fig. 1B) indicating the B cell repertoire rapidly expanded following infection giving rise to the neutralizing antibody diversity. Interestingly, only four mAbs (THSC20.HVTR04, THSC20.HVTR06, THSC20.HVTR11 and THSC20.HVTR26) were found to variably neutralize the Omicron variants when tested as pseudoviruses and authentic live viruses (Fig. 1C-D). THSC20.HVTR04 (IGVH3-30) though found to have lost its activity against BA.1 (7) presumably due to G446S and N440K mutations, it was found to potently neutralize BA.2, BA.4 (pseudovirus) and BA.5 (authentic live virus) with IC50 values of 0.29, 0.21 and 0.19 μg/mL respectively. On the other hand, THSC20.HVTR06 (IGVH7-4-1) could neutralize all the BA.1, BA.2 and BA.4/BA.5 variants but with low potencies, while THSC20.HVTR11 could neutralize BA.1 and BA.2 with comparable potency as THSC20.HVTR04, however failed to neutralize BA.4/BA5. THSC20.HVTR26 mAb that we reported to moderately neutralize BA.1 (7) was included in the experiment for comparison (Table S1). The N440K mutation which is common in Omicron BA.1, BA.2 and BA.4/BA.5 variants did not appear to affect the neutralization potential of THSC20.HVTR04 against BA.2 and BA.4/5, though likely have reduced its potency as noted earlier with SARS-CoV-2 wild type spike. Overall, we observed unique neutralization diversities conferred by the antibody repertoire developed in this particular individual through natural infection.

**Fig. 1.**
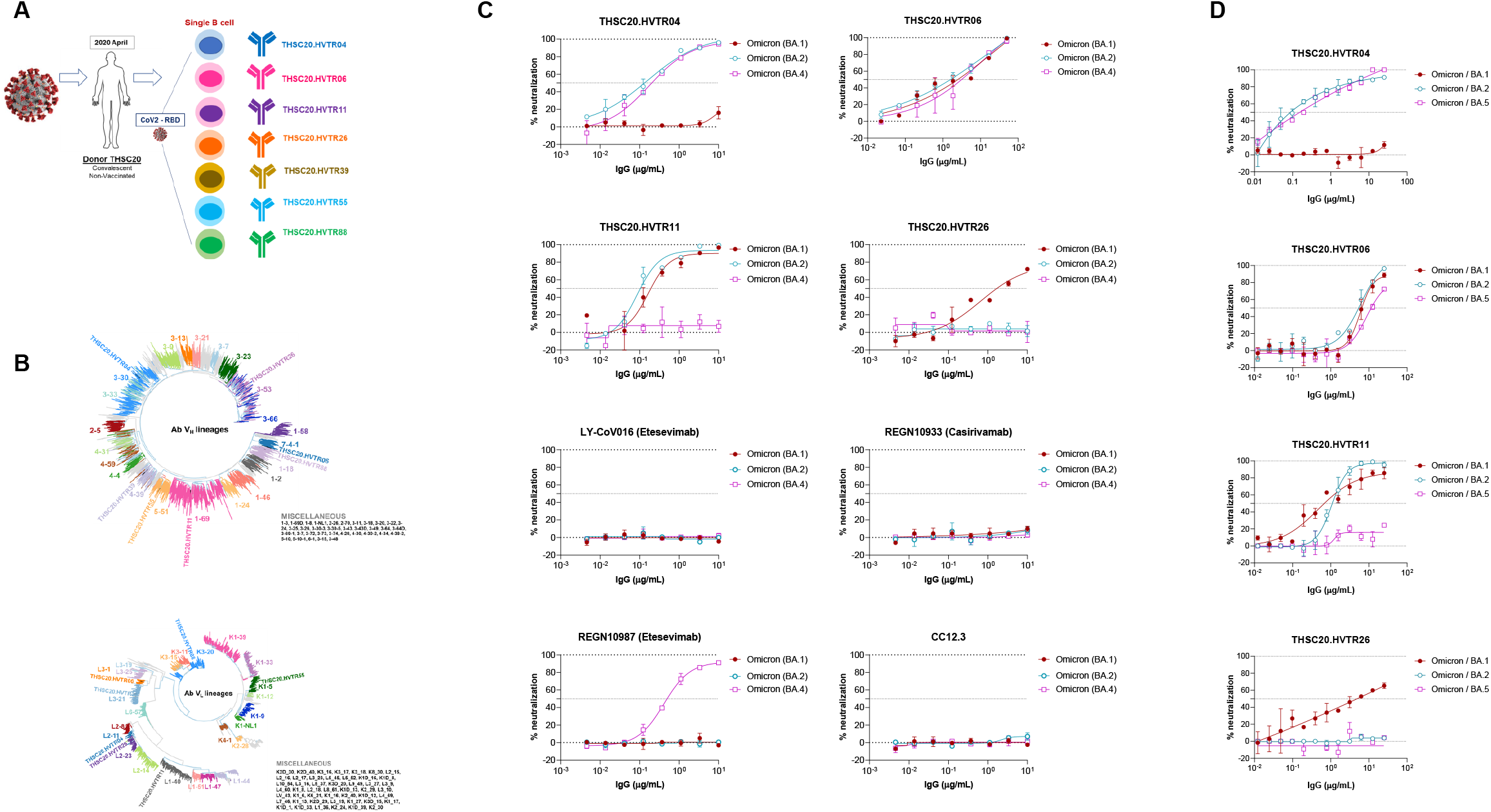
Diversity of the antibody repertoire developed in an unvaccinated individual infected with ancestral SARS-CoV-2. A. Representation of the seven different mAbs obtained from the C-03-0020 donor infected with the ancestral SARS-CoV-2 and those originated from distinct B cell germlines. B. Phylogenetic clustering of maximum likelihood trees of variable heavy and light chain immunoglobulin G (IgG) genes representing the 7 neutralizing mAbs isolated from the donor C-03-0020 along with 3441 variable heavy and light IgG chain genes representing human B cell germline obtained from CoVAbDAb database). SARS-CoV-2 ‘neutralizing’ monoclonal antibodies reported in CovAbDAb database (http://opig.stats.ox.ac.uk/webapps/covabdab/), obtained from human germlines were selected based on availability of both heavy and light chain sequences (8). Post filtration, a total of 3448 sequences were retained (including 7 mAb sequences from the present study). Variable heavy and light chain IgG amino acid sequences were aligned using MAFFT v7.453(10). Maximum likelihood trees were constructed with IQTree (V 1.6.12) using a custom Ab amino acid substitution model(11, 12). Robustness of the tree was assessed using 1000 ultrafast bootstrap replicates as well as 1000 aSH-LRT replicates. Trees were visualized and annotated using ggtree package in R(13). C. Dose-dependent neutralization of pseudoviruses expressing spikes of Omicron BA.1, BA.2 and BA.4 against THSC20.HVTR04, THSC20.HVTR06, THSC20.HVTR11 and THSC20.HVTR26 mAbs. Experiments were done in duplicates and repeated at least three times. D. Neutralization potency of THSC20.HVTR04, THSC20.HVTR06, THSC20.HVTR11 and THSC20.HVTR26 mAbs against live authentic Omicron BA.1, BA.2 and BA.5 assessed in focus reduction neutralization test (FRNT). Each point represents mean percent neutralization at a given dose of mAb plotted on the X-axis. Neutralization curves plotted using GraphPad Prism software (v8.1.2).

Next, we examined the magnitude and quality of neutralizing antibodies against Omicron variants developed in the same C-03-0020 donor and compared with antibody responses observed with individuals who were vaccinated post infection with ancestral SARS-CoV-2. The participants included in this study were members of DBT COVID-19 consortium cohort, organized by interdisciplinary research institutes and hospitals in the National Capital Region of India. The study protocol was approved by the Institute Ethics Committees of all participating institutions. Written informed consent was obtained from all the participants who contributed bio-specimens, and for the clinical information collected. As part of this case study, we first examined the magnitude of antibodies developed in the C-03-0020 donor, 20 weeks post receiving third vaccine dose of ChAdOx1nCoV-10 (Covishield^TM^) (Fig. 2A) for its extent to neutralize the Omicron BA.1, BA.2 and BA.4. Remarkably, antibodies obtained from the followed-up visit from this individual demonstrated neutralization of Omicron BA.1, BA.2 and BA.4 with significantly increased potency compared to its baseline convalescent antibodies by 16.06, 11.33 and 9.67 folds respectively (Fig.2B). Next, we compared the magnitude of antibody response developed post vaccination in this particular individual (C-03-0020) with that obtained from sixteen vaccine breakthrough Omicron infected individuals (Table S2) and from ten individuals who received two doses of ChAdOx1nCoV-10 (Covishield^TM^) but not reported to be infected post vaccination (Table S3) in neutralizing Omicron BA.1, BA.2 and BA.4 variants. Interestingly, the plasma antibodies obtained from C-03-0020 donor post receiving three doses of vaccines showed neutralization of the three Omicron variants examined with comparable magnitude to that observed with antibodies developed in Omicron BTI individuals which demonstrated most potent neutralization of the same viruses (Fig.2C; Table S1). This was observed in sharp contrast to that obtained from those who received full vaccine doses with no history of infection and re-infection as shown in Fig. 2C; Table S3). This indicates that the diversity of antibody responses mounted in this particular individual (C-03-0020) immediately post infection while collectively overcome the mutational landscape offered by the evolving SARS-CoV-2, development of cross neutralizing antibodies with higher magnitude following vaccine boosters observed was likely due to clonal expansion of the B cell linages developed in this individual through mutations within the antibody genes associated with affinity maturation as previously indicated elsewhere (9). To further investigate whether what we observed with this particular individual (C-03-0020) is unique or not, we examined the ability and magnitude of antibody responses developed in seven additional individuals who were infected with ancestral SARS-CoV-2 in April 2020 before and after receiving at least two doses of vaccines (Table S2). Although antibodies developed in these donors before vaccination varied in neutralizing Omicron variants, but the antibody/immune response mounted post vaccination, demonstrated neutralization of the same variants with increased potency (Fig. 2D) and which was comparable to that observed with C-03-0020 follow up plasma samples. Notably, we also made the following observations with the individuals previously infected with ancestral virus: (a) antibodies elicited post vaccination from one of them (C-10-0006) neutralized all the Omicron variants with significantly higher magnitude (potency) and (b) the ability to neutralize the same Omicron variants by plasma antibodies obtained post vaccinations from other donors varied - for example, antibodies elicited in donor C-03-0015 could neutralize all the Omicron examined in this study, while donor C-03-0008 could neutralize BA.1 and BA.2 but not BA.4 (Fig. 2D). Overall, these observations indicate that while vaccination post infection improved humoral immune responses, variability in their magnitude and potential to counter all Omicron variants was possibly be due to the differences in their ability to rapidly develop unique B cell repertoire post infection associated with mounting of robust polyclonal antibody responses. Individuals who were able to develop such strong polyclonal antibody responses post infection such as what we observed with C-03-0015 and C-03-0020 are more likely to be protected against emerging and future SARS-CoV-2 variants.

**Fig. 2.**
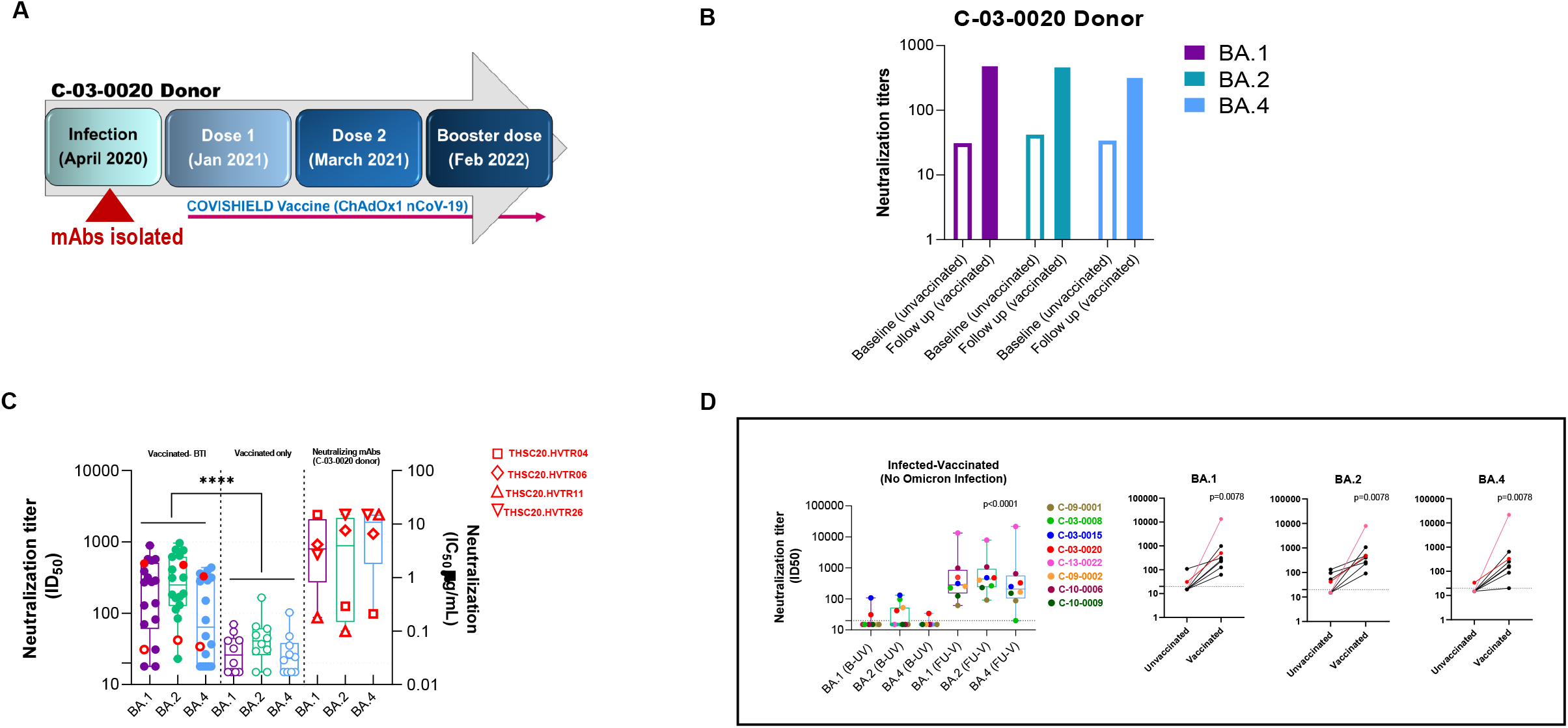
Neutralizing antibody response following vaccination in the C-03-0020 donor infected with the ancestral SARS-CoV-2. A. Schedule of vaccine (ChAdOx1nCoV19) doses and the intervals in between the doses received by the C-03-0020 donor; B. Neutralizing antibody responses of the plasma obtained from before and after full course (three doses) of vaccination against Omicron BA.1, BA.2 and BA.4 as assessed by pseudovirus neutralization assay; C. Comparison of the neutralization titer of antibodies developed in the C-03-0020 individual before and after vaccination with that developed in individuals with Omicron BTI and those who received two doses of vaccines (ChAdOx1nCoV19) with no history of infection. Red open and filled circles represent neutralization titer (ID50 doses) conferred by the plasma obtained from the C-03-0020 before and after receiving three doses of vaccine. ID50 refers to the reciprocal dilution of the heat-inactivated plasma samples that conferred 50% reduction in infection in a pseudovirus neutralization assay. The IC50 neutralization titers (IC50 values in μg/mL) of the four mAbs (THSC20.HVTR04, THSC20.HVTR06, THSC20.HVTR11 and THSC20.HVTR26) that showed variable neutralization potential against the Omicron variants are plotted as reference on the right. Significant difference in neutralizing antibody response developed between Omicron BTI and vaccinated but infected individuals was observed (p<0.0001) using Mann-Whitney statistical test D. Magnitude of neutralization antibody responses (ID50) of plasma obtained from 11 individuals infected with the ancestral SARS-CoV-2 (including C-03-0020 donor shown as red dot) before vaccination and the same post receiving two doses of vaccine by eight individuals (including C-03-0020 donor shown as red dot). B-UV refers to baseline unvaccinated and FU-V refers to follow up post full vaccination. Pre-vaccination versus post-vaccination neutralizing antibody titers of eight individuals infected with ancestral SARS-CoV-2 against Omicron BA.1, BA.2 and BA.4 as shown in the next three plots respectively, were found to be statistically significant (p < 0.01) with Wilcoxon matched-pairs signed-rank test.

## Acknowledgements

We are grateful to the study participants, our collaborators and partners for their valuable support. We also thank the members of the THSTI clinical team, data management and Biorepository for data records, sample processing and access to samples for the study. We are grateful to the members of the Institutional Ethics Committee for rigorous review and valuable input given on the study protocols and experimental designs. We thank Penny Moore and Jinal Bhiman, NICD, Johannesburg, South Africa to kindly provide us with the Omicron spike plasmid constructs for preparation of pseudoviruses, Jason McLellan, Texas, USA for kindly providing us with the SARS-CoV-2 RBD construct used for purification of RBD protein, IAVI-Neutralizing Antibody Center, The Scripps Research, La Jolla, California, USA for providing us with the REGN-10933 (Casirivimab), REGN-10987 (Imdevimab) and LY-CoV016 (Etesevimab) monoclonal antibodies. We finally thank all the members of our laboratory, THSTI and Advanced Technology Platform Center (ATPC) of the Regional Center of Biotechnology for antibody sequencing and access to BLI support for their support. This study was supported by funding from the Bill and Melinda Gates Foundation, Seattle, USA (INV-030592), Indian Council of Medical Research (CTU/Cohort study/17/10/22/2021/ECD), Department of Biotechnology (BT/PR40401/BIOBANK/03/2020), Research Council of Norway (Project ID: 285136), JB is supported by the DBT-Wellcome Trust India Alliance Team Science Grant (IA/TSG/19/1/600019). Dr PKG is supported by the J C Bose fellowship from SERB.

## Supplementary Information

**Table S1**. Neutralization breadth and potency of the mAbs isolated from infected but not vaccinated C-03-0020 donor.

**Table S2**. History of vaccination prior to infection with Omicron variant (BTI) and neutralization antibody responses post infection against Omicron BA.1, BA.2 and BA.4 variants

**Table S3.** Vaccination history of individuals previously infected with ancestral SARS-CoV-2 and neutralization antibody responses against Omicron variants post vaccination.

**Fig. S1. Characterization of newly isolated neutralizing mAbs**. A. Nucleotide sequences of the CDRH3 and CDRL3 regions of variable heavy and light IgG chains of THSC20.HVTR11 and THSC20.HVTR55 and their B cell allelic origins as determined by using the international ImMunoGeneTics information system database (http://www.imgt.org). B. Effect of L452R substitution on the ability of THSC20.HVTR11 to neutralize SARS-CoV-2 in a pseudovirus neutralization assay. Pseudoviruses expressing spikes of Kappa (B.1.617.1), Delta (B.1.6517.2) and Omicron BA.4 which naturally contains L452R were included in the same assay. The neutralization assay was done in duplicates and was repeated at least three times. C. Binding avidity of the newly isolated THSC20.HVTR11 and THSC20.HVTR55 to SARS-CoV-2 by ELISA. D. Binding affinities of THSC20.HVTR11 and THSC20.HVTR55 to the SARS-CoV-2 (Wuhan) receptor binding domain (RBD) protein by BLI-Octet. Binding kinetics of THSC20.HTR11 and THSC20.HVTR55 with SARS-CoV-2 RBD using BLI-octet analysis. His-tagged SARS-CoV-2 RBD was immobilized on NTA biosensors and binding kinetics of the IgGs to RBD was assessed using four different concentrations starting with 22.3 nM followed by 3-fold dilutions till 0.8 nM. Association and dissociation of the mAbs to the RBD bound biosensors were assessed for 500 sec each. Data shown is reference - subtracted and analyzed using Octet data analysis software v11.1 (Forte Bio Inc.). Global curve fitting using 1: 1 binding model determined K_on_, K_off_ and K_D_ values shown below the graphs. E. Epitope binning assay. THSC20.HVTR11 and THSC20.HVTR55 were evaluated for epitope competition using BLI. All the incubation steps for binning experiments were performed in 1x PBS. 50-100 nM of his-tagged RBD protein antigens were loaded on Ni-NTA biosensors to achieve 0.9 to 1.3 nm of wavelength shift and then washed. Saturating concentration of mAbs (100μg/ml) was added for 10 min and competing mAbs at concentrations of 25 μg/ml were then added for 5 min in order to measure binding in the presence of saturating antibodies.

